# Enteric nervous system degeneration in human and murine CLN3 disease, is ameliorated by gene therapy in mice

**DOI:** 10.1101/2025.01.29.635518

**Authors:** Ewa A. Ziółkowska, Letitia L. Williams, Matthew J. Jansen, Sophie H Wang, Elizabeth M. Eultgen, Jaiprakash Sharma, Marco Sardiello, Rebecca P. Bradley, Ineka T. Whiteman, Mark S. Sands, Robert O. Heuckeroth, Jonathan D. Cooper

## Abstract

**Background and aims:** Severe gastrointestinal (GI) symptoms occur in people with CLN3 disease, a neurodegenerative disorder. If left untreated these GI symptoms compromise life quality and may contribute to death. We hypothesized GI symptoms in CLN3 disease are at least partially due to neurodegeneration in the enteric nervous system (ENS), the master regulator of bowel function.

**Methods:** We examined the integrity of the ENS in human CLN3 autopsy small bowel and colon, and in CLN3 deficient (*Cln3^Δex^*^7^*^/^*^8^) mice. We performed detailed immunohistological analyses of enteric neurons and glia and assessed bowel transit times at multiple disease stages. We then tested the therapeutic potential of neonatal intravenous gene therapy (AAV9-hCLN3) to prevent bowel phenotypes in *Cln3^Δex^*^7^*^/^*^8^ mice.

**Results:** Human CLN3 bowel displayed a profound loss of enteric neurons and their neurites, with pathological effects upon enteric glia. *Cln3^Δex^*^7^*^/^*^8^ mice had normal appearing ENS at 1 month of age, but then experienced progressive loss of both enteric neurons and glia accompanied by marked bowel distention, resembling the human CLN3 phenotype. Degenerative changes in *Cln3^Δex^*^7^*^/^*^8^ mouse enteric neurons and glia were largely prevented by systemic neonatal delivery of AAV9-hCLN3 gene therapy, preventing bowel distention at disease endstage.

**Conclusions:** Our findings demonstrate that CLN3 deficiency profoundly damages enteric neurons and glia in both murine and human CLN3 disease, contributing to GI dysfunction. This study provides preclinical evidence that systemic gene therapy may effectively treat multiple aspects of bowel pathology, expanding the therapeutic landscape beyond the CNS.

What you need to know:

**Background and Context:** Significant gastrointestinal (GI) symptoms are evident in many pediatric neurological conditions. We hypothesized that, in addition to central nervous system (CNS) effects, defects in the enteric nervous system (ENS) may underlie these GI symptoms in some neurodegenerative diseases. Revealing such defects would open up new opportunities for treating these life-limiting and debilitating symptoms.

**New Findings:** The enteric nervous system is significantly impacted in human CLN3 disease, a feature that is recapitulated in CLN3 mice. Progressive enteric neurodegeneration in these mice follows a similar time course to neuron loss in the brain, resulting in severe bowel distention.

Nevertheless, bowel distention and the majority of the pathology within the enteric nervous system can be mitigated via neonatal gene therapy.

**Limitations:** Our human data will need to be replicated in larger numbers of CLN3 cases, and methods will need to be developed to treat the human bowel, avoiding the risk of liver tumors.

**Impact:** These results reveal that a neurodegenerative disease previously thought to primarily affect the CNS, damages the bowel’s enteric nervous system and that ENS degeneration can be prevented in mice by gene therapy. These data provide a new perspective on this pediatric disorder and may have relevance to other pediatric neurologic diseases.

**Lay Summary:** The progressive loss of neurons in CLN3 disease is not confined to the brain but also occurs in the bowel enteric nervous system, contributing directly to GI dysfunction.

Neurodegeneration in the enteric nervous system can be prevented by treating the bowel with gene therapy.

## Introduction

Central nervous system (CNS) neurodegenerative diseases that impair cognitive and motor function are often accompanied by bowel-associated symptoms including constipation, vomiting, gastro-esophageal reflux, abdominal distension, abdominal pain and malnutrition. While these symptoms may be attributed to CNS dysfunction or medicine side effects, some mechanisms that damage the CNS might also damage the enteric nervous system (ENS). The ENS is the intrinsic nervous system of the bowel that senses local mechanical and chemical stimuli and then coordinates bowel muscle contraction and relaxation to facilitate nutrient absorption and eliminate waste^1–6^. The ENS also controls local blood flow, influences immune cell activity, and regulates epithelial function to maintain fluid and electrolyte balance and protect from infection. To complete these tasks, the ENS has about as many neurons as the spinal cord, ∼20 enteric neuron subtypes and ∼8 types of enteric glia^1–9^. Not surprisingly, ENS defects cause constipation, vomiting, malnutrition, abdominal distension, abdominal pain, and may predispose to bowel inflammation. While inflammation and malnutrition reduce quality of life, they also exacerbate some CNS neurodegenerative diseases^10–15^.

We explored these issues in CLN3 disease (also called juvenile neuronal ceroid lipofuscinosis (JNCL)), a pediatric neurodegenerative disorder in which GI symptoms become prominent as the disease progresses, although very little is published regarding these well recognized symptoms^16^. CLN3 disease is the most common of a larger group of disorders termed neuronal ceroid lipofuscinosis (NCLs)^16–21^, more commonly known as Batten disease. Mutations in the *CLN3* gene cause deficiency of the lysosomal CLN3 transmembrane protein, which is present in all cells^22^, leading to profound CNS neurodegeneration^23–31^ through mechanisms that remain poorly understood. It has been suggested that glial cells may play an active role in CLN3 pathogenesis, directly contributing to neuron loss^32–35^. In addition to its obvious neurological effects, CLN3 disease causes a range of symptoms in the rest of the body, including effects upon cardiac, autonomic and neuromuscular systems^16–18,31^. Such symptoms may occur secondary to degeneration within the CLN3 CNS, but we hypothesized that bowel dysfunction could also be caused by degeneration of neurons and glia of the ENS that are negatively affected by CLN3 deficiency.

To determine if the ENS is compromised in CLN3 disease, we assessed autopsied human bowel and found evidence of a profound degeneration of enteric neurons and glia. We then used the *Cln3^Δex^*^7^*^/8^*mouse model^36^ to define the onset and progression of ENS pathology and its functional consequences. These analyses revealed severe distention of the bowel that worsened with age, atrophy of bowel smooth muscle and progressive loss of enteric neurons and glia in all bowel regions of *Cln3^Δex7/8^* mice. To prevent this ENS degeneration, we employed gene therapy^17–19,37^, a potential cure for this monogenic disorder if CLN3 protein could be delivered to the right cells. Because CNS targeted gene therapy^38–40^ would be unlikely to efficiently treat the ENS manifestations of CLN3 deficiency, we tested whether neonatal intravenous (IV) delivery of an adeno-associated viral vector (AAV9) expressing human CLN3 to *Cln3^Δex7/8^* mice would prevent CLN3 disease effects in the bowel. This AAV9 vector targets enteric neurons, but not glia or bowel smooth muscle^41^. However, neonatal AAV9-mediated gene therapy reduced bowel distention and largely prevented the loss of enteric neurons and glia at disease endstage. Taken together our data reveal novel effects of CLN3 deficiency upon the bowel and suggest these disease manifestations may be preventable by bowel-directed gene therapy. These findings have potential implications for developing effective therapies for this and other similar disorders.

## MATERIALS AND METHODS

### Human bowel samples

Human CLN3 bowel samples were obtained at autopsy with fully informed consent from an individual confirmed to be homozygous for the common 1.02kb deletion in the *CLN3 g*ene^22^. Bowel was shared with investigators as a de-identified sample. This individual had a typical CLN3 disease onset and died at 21 years of age. The use of this de-identified sample at Washington University School of Medicine in St. Louis was Institutional Review Board exempt. The control 18-year-old human colon was an organ donor, while the 17-year-old human ileum was from a de-identified subject who had surgery at Children’s Hospital of Philadelphia (CHOP) with approval of CHOP’s IRB (IRB 13-010357). Organ donor colon was from the Gift of Life Donor Program (also Institutional Review Board exempt).

## Mice

*Cln3^Δex7/8^* mice^36^, and wild type (WT) mice were maintained separately on a congenic C57Bl/6J background at Washington University School of Medicine. Mice were provided a standard mouse chow (Purina Rodent Diet 5053) and water *ad libitum* under a 12hr light/dark cycle. Numbers analyzed are detailed in Figures, but all studies used balanced numbers of males and females.

These studies follow ARRIVE guidelines and were performed under protocols 2018-0215, 21- 0292 and 24-0232 approved by the Institutional Animal Care and Use Committee (IACUC) at Washington University School of Medicine in St. Louis, MO.

## Bowel transit studies

*Whole bowel transit*: Mice fasted overnight were gavage-fed with 6% carmine red dye (300 µL) Sigma-Aldrich) in distilled water containing 0.5% methylcellulose (MilliporeSigma) and placed in individual cages with white paper covering cage bottoms. Passage of red stool was evaluated at 10 min intervals. Each mouse was tested three times with ∼3 days between tests, as previously described^41–44^.

*FITC-dextran transit*: Mice fasted overnight were gavage-fed with 100 μL FITC-Dextran (10 mg/mL, 70,000 MW; MilliporeSigma) in distilled water containing 2% methylcellulose (MilliporeSigma). After 2 hours, bowel from isoflurane-euthanized mice was divided into 15 segments (small intestine 1–10, proximal and distal cecum, colon 1–3). Segment contents were suspended in 100 µL 1X phosphate buffered saline, vortexed 15 s, and centrifuged (4000 rpm, 10 min). Supernatant fluorescence was measured (excitation 485 nm, emission at 525 nm) on a plate reader. Weighted geometric mean (sum of (segment number × FITC fluorescence in that segment)/total FITC fluorescence) was calculated as described^41–44^.

*Colonic bead expulsion*: Mice were anesthetized with 2% isoflurane (1.5 L/min). A glass bead (3 mm, MilliporeSigma) lubricated with sunflower seed oil (MilliporeSigma) was inserted 2 cm into the colon using a custom-made 3 mm diameter rounded glass rod. Anesthesia was stopped and bead expulsion time was recorded. Assay was repeated 3 times per mouse with >48 hours between procedures^41–44^.

## Human bowel wholemount immunostaining and imaging

Wholemount specimens of human bowel were processed via a previously published clearing and immunostaining protocol^45–47^. Briefly, following trimming of visceral fat and fixation, 5 mm x 5 mm full thickness pieces of bowel were washed in PBS, permeabilized with 100% methanol, treated with Dent’s bleach and placed in blocking solution (3 days on a shaker at 37°C). This was followed by incubation in primary antibody (Supplementary Table 1) (14 days at 37°C) on a shaker. Tissue was then washed in PBS followed by incubation in secondary antibody (Supplementary Table 1) (3 days at 37°C) on a shaker, dehydrated in serial methanol dilutions, and cleared with benzyl alcohol-benzyl benzoate (BABB) (1:2) until tissue was translucent (∼24 hours). Tissue was transferred to ethyl cinnamate and mounted on glass slides in ethyl cinnamate for imaging via confocal microscopy within 5 days of tissue clearing^45–47^.

Cleared human bowel tissue was imaged on a Zeiss LSM 980 confocal microscope (10x/0.3 and 20x/0.8 Plan-Apochromat objectives, and *Zen* 3.5.093.00003 software (Zeiss, Oberkochen, Germany). *Z*-axis intervals were 4 mm (10x objective) or 1 mm (20x objective). For each segment imaged, confocal Z-stacks were obtained, and 10x objective imaging used tile scan (5% overlap) and stitch features to visualize large regions of full-thickness tissue. Multichannel images were acquired sequentially using laser-scanning operated under multitrack. Excitation/ long-pass emission filters were Alexa Fluor 594, and Alexa Fluor 647. Excitation/ long-pass emission filter Alexa Fluor 488 was used to capture autofluorescent storage material (ASFM) autofluorescence^45–47^.

## Wholemount bowel histology in mice

Mice were anesthetized (2% isoflurane), then transcardially perfused with PBS and decapitated. The first and last 8 cm of small intestine and entire colon were cut into 2 cm lengths in cold 50 mM Tris buffered saline (TBS, pH=7.6), opened along mesenteric border, pinned to Sylgard^TM^ 184 Silicone Elastomer (Dow Corning) using stainless steel insect pins, and fixed in fresh 4% paraformaldehyde (35 minutes colon, 45 minutes ileum, 1 hour duodenum)^32^. Fixed tissue was stored in TBS/0.1% sodium azide (4°C) before staining for neuronal or glial markers^41–44^. Muscle layers (1 cm lengths) separated from mucosa and submucosa were blocked (1 hour) in TBS containing 4% Triton X-100 (TBST), 15% normal goat serum (NGS) (Jackson Immuno-Research Laboratories), then incubated in primary antibody (Supplemental Table 1) at 4°C overnight, washed (3x, TBS), and incubated in secondary antibodies (Supplemental Table 1) in 10% goat serum, 4% TBST for 2 hours. Bowel was mounted on Colorfrost plus (Thermo Fisher Scientific) slides, dried briefly before incubation in 1x solution TrueBlack lipofuscin autofluoresence quencher (Biotium) in 70% ethanol (5 min) and rinsing (1xTBS). Slides were coverslipped in Fluoromount-G mounting medium containing DAPI (Southern Biotech)^41^.

## Quantitative analysis of immunostained mouse bowel

Unless otherwise specified, 10 systematically sampled regularly spaced 20× fields (each 0.4225 mm^2^) per sample were analyzed^41^. Multi-channel images were collected on a Zeiss AxioImager Z1 microscope using StereoInvestigator (MBF Bioscience) software and exported to ImageJ (NIH). Myenteric neurons (HuC/D+ cells) and enteric glia (S100B+ cells) were manually counted in each field and averaged to determine cells/mm^2^ ^41^. All analyses were made blind to genotype or treatment.

## Intravenous gene therapy

Injections employed a AAV9 virus expressing human CLN3 (hCLN3). This is essentially the same virus used in our CNS studies in CLN1 disease mice^48–54^ and described previously in detail^48^, but expressing the hCLN3 sequence instead of the PPT1 sequence. Virus was packaged at University of North Carolina Vector Core. Virus injections (1.5 x 10^11^ vg/mouse) were made into the superficial temporal vein of hypothermia-anesthetized neonatal (P1) mice^41^. Neonatally injected mice were returned to their mothers, raised until weaning, and were aged until predicted disease endstage, at which point most analyses were conducted.

## Statistical analyses

Statistical analyses were performed using GraphPad Prism version 9.1.0 for MacOS (GraphPad Software, San Diego, CA), assuming a normal distribution of data. A one-way ANOVA with a post-hoc Bonferroni correction was used for comparison between three groups or more.

Unpaired t-tests were used for comparisons between two groups. A p-value of ≤0.05 was considered significant. In all experiments the relevant control groups were included as indicated in each figure caption, along with the number of animals used in each experiment, with individual animals being the experimental unit. No randomization was performed and no data excluded.

## RESULTS

### Human CLN3 disease small bowel and colon displays ENS pathology

To examine the ENS in human CLN3 bowel, we obtained small bowel (ileum) and colon at autopsy from a 21-year-old female with genetically confirmed CLN3 disease. Full thickness ileum and colon were pinned flat, fixed, stained with antibodies, cleared and then imaged as a wholemount preparation (without sectioning) using confocal microscopy to generate 3- dimensional images of the ENS^41,45–47^ (**Figure 1**). To compare to this CLN3 disease bowel, we similarly processed human ileum from a 17-year-old male undergoing surgery and colon from an 18-year-old female organ donor. Human bowel wholemount preparations were stained with antibodies to HuC/D (all nerve cell bodies), Tuj1 (to identify neurites), and SOX10 (to identify enteric glia). In these same regions we imaged autofluorescent storage material (AFSM), a hallmark of NCL pathology^16–21^. These analyses revealed marked differences in the density of enteric neurons within ganglia (**Figure 1A**) and regions where ganglia appeared to be missing, although these changes were highly variable between ganglia. We also detected a pronounced accumulation of AFSM in HuC/D+ enteric neurons from the CLN3 disease ileum (**Figure 1B**). At higher magnification, changes in the morphology of HuC/D+ neurons were evident in the CLN3 disease ileum (**Figure 1A, B**), with fewer large neurons compared to the control organ donor ileum (**Figure 1A, B**). There were also markedly fewer SOX10+ glia in the human CLN3 disease ileum than in control ileum, with many SOX10+ HuC/D+ double positive cells in this human CLN3 disease ileum that were uncommon in control bowel (**Figure 1B**). Similar effects were evident for colon enteric neurons in this CLN3 bowel, with fewer large neurons, a dramatic loss of Tuj1-positive neurites, and marked AFSM within HuC/D+ neurons compared with the control organ donor colon (**Figure 1C**). These data suggest CLN3 disease damages enteric neurons and glia in humans and prompted us to investigate whether similar phenotypes are evident in a mouse model of CLN3 disease.

**Figure 1.**
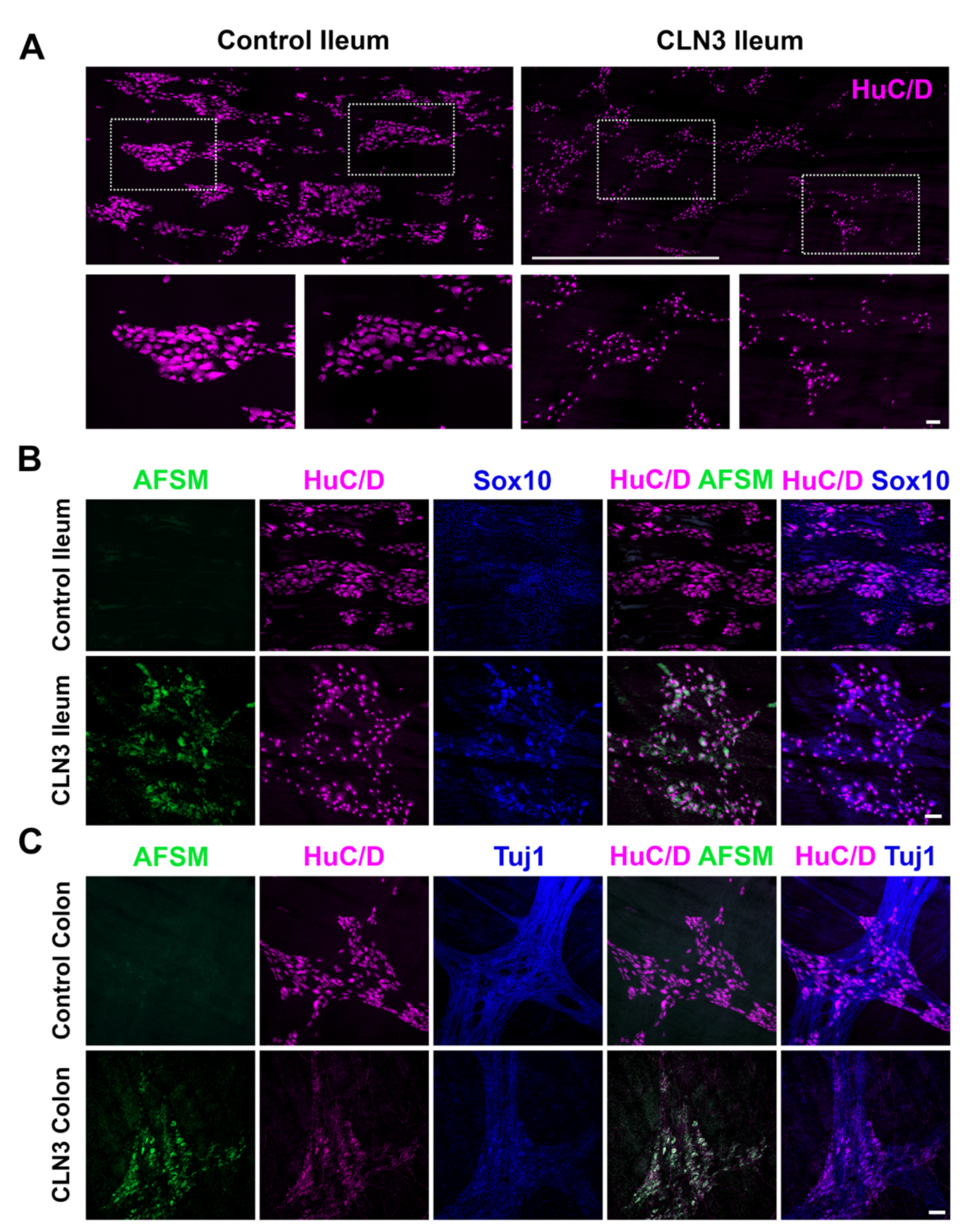
Enteric nervous system pathology in human bowel from a young adult with CLN3 disease. (**A**) Human ileum from a previously healthy 17-year-old male (left) and ileum from a 21-year-old female young adult who died from CLN3 disease (right) was stained with antibodies to pan-neuronal protein HuC/D (magenta), cleared, and imaged by confocal microscopy from mucosa to serosa with a 10X objective. Large images show stitched confocal Z-stacks of layers containing myenteric plexus. Scale bar 1000µm. Lower panels are magnifications of the regions enclosed in boxes from large tiled images in the top row. Scale bar 100µm. (**B**) Ileum from the same healthy 17-year-old and the same young adult with CLN3 disease was stained with antibodies to HuC/D (magenta) and SOX10 (blue), cleared and imaged by confocal microscopy from mucosa to serosa with a 10X objective. Images show autofluorescent storage material (AFSM, green), HuC/D and SOX10 signals in a single slice from the confocal Z-stack in the layer containing myenteric plexus. Scale bar 100µm. (**C**) Colon from a previously healthy 18-year-old female organ donor and the same young adult with CLN3 disease was stained with antibodies to HuC/D (magenta) and Tuj1 (blue), cleared and imaged by confocal microscopy from mucosa to serosa with a 10X objective. Images show AFSM, HuC/D and Tuj1 signals a single slice from the confocal Z-stack in the layer containing myenteric plexus. Scale bar 100µm.

## Bowel distention and transit phenotypes in *Cln3^Δex7/8^* mice

*Cln3^Δex7/8^* mice bear a deletion in exon 7 and 8 of the *Cln3* gene^36^ that is equivalent to the common 1.02kB deletion present in 85% of alleles in human CLN3 disease^22^. We previously characterized the onset and progression of *Cln3^Δex7/8^* mouse CNS disease phenotypes^26–31^ and conducted pre-clinical studies designed to treat these disease manifestations^38,55–61^. To begin exploring the impact of CLN3 deficiency on gastrointestinal function we examined the bowel of *Cln3^Δex7/8^* mice at disease endstage (18 months of age). At this age, *Cln3^Δex7/8^*bowel appeared markedly distended, with stool filled proximal colon, cecum and distal small bowel **(Figure 2A)**. This phenotype suggested bowel dysmotility in *Cln3^Δex7/8^* mice, which we investigated by assessing bowel transit rates^41–44^. Whole bowel transit as assessed by Carmine red revealed *Cln3^Δex7/8^* mice and age-matched WT controls were not different at any age, although mean transit times were consistently longer in *Cln3^Δex7/8^* mice at each age **(Figure 2B)**, and slowed with increasing age in both *Cln3^Δex7/8^* and WT mice. The distribution of FITC-labelled dextran 2 hours post gavage^41–44^ was also statistically the same in WT and *Cln3^Δex7/8^* mice, although mean FITC-dextran levels trended higher in distal small bowel (35% increase) and lower in cecum (43% decrease) and proximal colon (56% decrease) **(Figure 2C)** of *Cln3^Δex7/8^*mice. Weighted geometric means of the FITC-dextran distribution were also not statistically different between groups (data not shown). To examine the impact of CLN3 deficiency on distal colon motility, we evaluated colonic bead expulsion times^41–44^. While mean bead expulsion time was 7.78% longer in *Cln3^Δex7/8^* mice compared to WT, this difference was not statistically significant **(Figure 2D)**.

**Figure 2.**
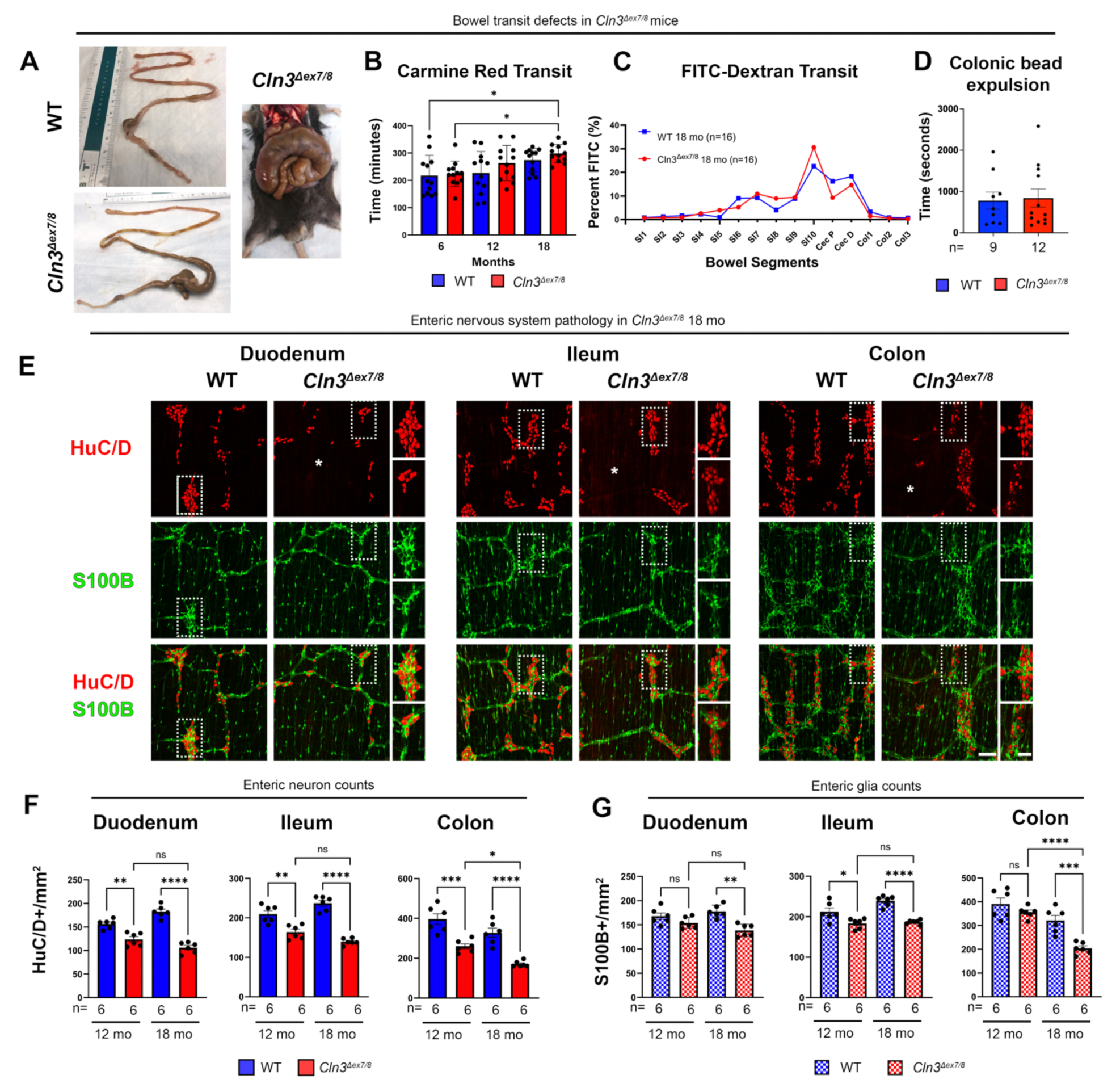
Enteric nervous system pathology in *Cln3^Δex7/8^* mice. **(A)** *Cln3^Δex7/8^* mice display distention of the distal small intestine, cecum and proximal colon by fecal material at disease end-stage. Images from *Cln3^Δex7/8^* mice show an extreme example. **(B,C)** Bowel transit was assessed by measuring the time for gavaged Carmine red to appear in stool **(B)** or determining the amount of FITC-conjugated dextrans present in individual bowel segments 2 hours after gavage **(C)** (SI1-SI10 = numbered segments of small intestine; Cec P=proximal cecum; Cec D=distal cecum; Col1-Col3=numbered segments of colon). **(D)** Time taken to expel a glass bead from the colon was measured at disease endstage. **(E)** Whole-mount bowel preparations immunostained for pan-neuronal marker HuC/D (red) and glial marker S100B (green) reveal the loss of myenteric plexus neurons and glia, and changes in neuron morphology in the duodenum, ileum and mid-colon of disease endstage *Cln3^Δex7/8^* mice vs. age matched wildtype (WT) controls at 18 months (mo). Scale bar 200µm, 100µm in magnified inserts, respectively. Asterix (*) indicates regions of near complete enteric neuron loss**. (F, G)** The density of HuC/D+ neurons and S100B+ enteric glia was measured in these mice at 12-month-old and at endstage (18- month-old). Data ± SEM (B, D, F, G). One-way ANOVA with a post-hoc Bonferroni correction **(B, C, F, G)**, unpaired t-test **(D)**. * p≤0.05, ** p≤0.01, *** p≤0.001, **** p≤0.0001, ns = not significant.

Together, these data suggest *Cln3^Δex7/8^* mice have normal bowel transit times that slow with age like WT, but very dilated bowel suggests bowel neuromuscular function is not normal. Consistent with these observations, we recently reported *Cln3^Δex7/8^* mice^62^ have bowel smooth muscle atrophy which may contribute to bowel distension along with CLN3 disease effects upon the ENS that is also evident in human CLN3 disease **(Figure 1)**.

### Progressive loss of enteric neurons in *Cln3^Δex7/8^* mice

To evaluate ENS myenteric neuron and glial cell density in small bowel and colon, we stained bowel wholemount preparations from *Cln3^Δex7/8^* mice and age-matched WT controls at different stages of disease progression for the neuronal marker HuC/D^41^ and the intestinal glial marker S100B^41,63^. We first performed these analyses at 1 month (presymptomatic disease stage), revealing a similar ENS appearance in mice of both genotypes **(Figure S1)**. Quantitative analysis confirmed no difference in the density of HuC/D positive neurons between WT and *Cln3^Δex7/8^* mice in any bowel region at 1 month **(Figure S1)**, suggesting normal ENS development.

To investigate whether apparently normal ENS development in *Cln3^Δex7/8^* mice was followed by progressive degeneration of enteric neurons, we next analyzed myenteric neuron density at disease midstage (12 months) and endstage (18 months) after HuC/D immunostaining.

Quantitative analyses showed significant reductions in the density of myenteric neurons in all bowel regions of 12-month-old *Cln3^Δex7/8^* mice compared to WT controls **(Figure 2E, 2F. S2A)** that became more pronounced with age. For example, myenteric neuron density in *Cln3^Δex7/8^* duodenum, ileum, and colon was 21%, 22% and 35% lower than WT, respectively at 12 months of age. By disease endstage (18 months), *Cln3^Δex7/8^* mice had only 42%, 41% and 48% of WT myenteric neuron density in duodenum, ileum and colon, suggesting progressive myenteric neuron loss **(Figure 2F**, **Table 1)**. Enteric neuron loss was not evenly distributed in *Cln3^Δex7/8^* myenteric plexus but occurred in patches with marked neuron loss adjacent to areas where neurons survived, and other areas where HuC/D-positive neurons appeared shrunken, similar to findings in human CLN3 bowel at autopsy **(Figure 1)**. These changes in enteric neuron density in both small bowel and colon were accompanied by a corresponding qualitative decrease in staining for the neuron-specific microtubule marker TuJ1 in the *Cln3^Δex7/8^* mice, that also became more pronounced with age **(Figure S2B)**. These data clearly indicate that there is a significant impact of CLN3 disease upon the integrity of enteric neurons in *Cln3^Δex7/8^* mice, resembling the pathology evident in human CLN3 bowel at autopsy.

**Table 1.** *Cln3^Δex7/8^* Histology Data at 12 months

| HuC/D | WT<br>neurons/mm<sup>2</sup> | *Cln3*<sup>Δex7/8</sup><br>neurons/mm<sup>2</sup> | Percent (%) Loss | p-value<br>(WT vs.<br>*Cln3*<sup>Δex7/8</sup>) |
| --- | --- | --- | --- | --- |
| Duodenum | 156.41±4.05 | 123.71±5.74 | 20.91% | p=0.0009 |
| Ileum | 209.63±9.00 | 164.22±7.28 | 21.66% | p=0.0029 |
| Colon | 396.73±27.24 | 260.12±12.61 | 34.43% | p=0.0011 |

| S100B | WT<br>S100B+<br>Ganglionic glia/<br>mm<sup>2</sup> | *Cln3*<sup>Δex7/8</sup> S100B+<br>Ganglionic glia/<br>mm<sup>2</sup> | Percent (%) Loss | p-value<br>(WT vs.<br>*Cln3*<sup>Δex7/8</sup>) |
| --- | --- | --- | --- | --- |
| Duodenum | 167.73±6.78 | 154.08±4.76 | 8.14% | 0.1305 |
| Ileum | 212.03±9.53 | 183.67±5.13 | 13.38% | 0.0255 |
| Colon | 391.01±24.92 | 355.82±9.78 | 9.00% | 0.2180 |

***Cln3*<sup>Δex7/8</sup> Histology Data at 18 months**
| HuC/D | WT<br>neurons/mm<sup>2</sup> | *Cln3*<sup>Δex7/8</sup><br>neurons/mm<sup>2</sup> | Percent (%) Loss | p-value<br>(WT vs.<br>*Cln3*<sup>Δex7/8</sup>) |
| --- | --- | --- | --- | --- |
| Duodenum | 182.60±5.31 | 106.15±4.69 | 41.87% | p<0.0001 |
| Ileum | 237.04±7.41 | 139.68±3.33 | 41.07% | p<0.0001 |
| Colon | 327.46±23.32 | 170.81±6.07 | 47.84% | p<0.0001 |

| S100B | WT<br>S100B+<br>Ganglionic glia/<br>mm<sup>2</sup> | *Cln3*<sup>Δex7/8</sup> S100B+<br>Ganglionic glia/<br>mm<sup>2</sup> | Percent (%) Loss | p-value<br>(WT vs.<br>*Cln3*<sup>Δex7/8</sup>) |
| --- | --- | --- | --- | --- |
| Duodenum | 177.75±5.28 | 138.66±5.25 | 21.99% | 0.0004 |
| Ileum | 239.25±5.26 | 186.71±2.24 | 21.96% | <0.0001 |
| Colon | 321.18±21.39 | 203.87±10.02 | 36.53% | 0.0006 |

## Effects of CLN3 deficiency upon enteric glia

ENS function also depends on enteric glia^2-4^, which may influence enteric neuron survival and function. To investigate the impact of CLN3 disease upon enteric glia we stained bowel wholemounts from *Cln3^Δex7/8^*mice for S100B and GFAP, two markers that are expressed by partially overlapping populations of enteric glia^63^. These analyses revealed an uneven and patchy distribution of GFAP immunoreactivity in the bowel of *Cln3^Δex7/8^* mice **(Figure S2B),** with areas of both increased GFAP immunoreactivity in some bowel regions, and other areas where GFAP immunoreactivity was markedly decreased. Areas with decreased GFAP immunoreactivity appeared to correlate with regions where decreased HuC/D neuron density was most pronounced. To determine whether *Cln3^Δex7/8^* mice have fewer enteric glia than WT, we measured the density of S100B positive enteric glia in different bowel regions. These analyses revealed significant reductions in the density of S100B positive enteric glia that only became apparent at disease endstage, with a 22% reduction in S100B glial density in duodenum and ileum of 18-month-old *Cln3^Δex7/8^* mice, and 37% reduction in their colon **(Figure 2G**, **Table 1).**

## AAV9-mediated gene therapy preserves ENS structure

Theoretically CLN3 deficiency is amenable to gene therapy^19,37^, and CNS-directed gene therapy has positive treatment effects upon the brains of *Cln3^Δex7/8^* mice^38–40^. To apply a gene therapy strategy to the bowel, we intravenously delivered 1.5 x 10^11^ vg/mouse of AAV9.hCLN3 to neonatal *Cln3^Δex7/8^* mice and subsequently investigated the impact of this therapy upon the changes we had defined in ENS structure and bowel function.

Compared to untreated *Cln3^Δex7/8^* mice, our analyses showed reduced mean Carmine red transit time in AAV9.hCLN3 treated *Cln3^Δex7/8^* mice at both 12 and 18 months but differences were not statistically significant **(Figure 3A)**. We next analyzed anatomic integrity of the ENS in wholemount bowel preparations from 18-month-old AAV9.hCLN3 treated *Cln3^Δex7/8^* mice **(Figure 3B)**. These analyses revealed a statistically significant protective effect of AAV9.hCLN3 treatment upon both HuC/D+ myenteric neurons and S100B+ myenteric glia **(Figure 3B, 3C)**.

**Figure 3.**
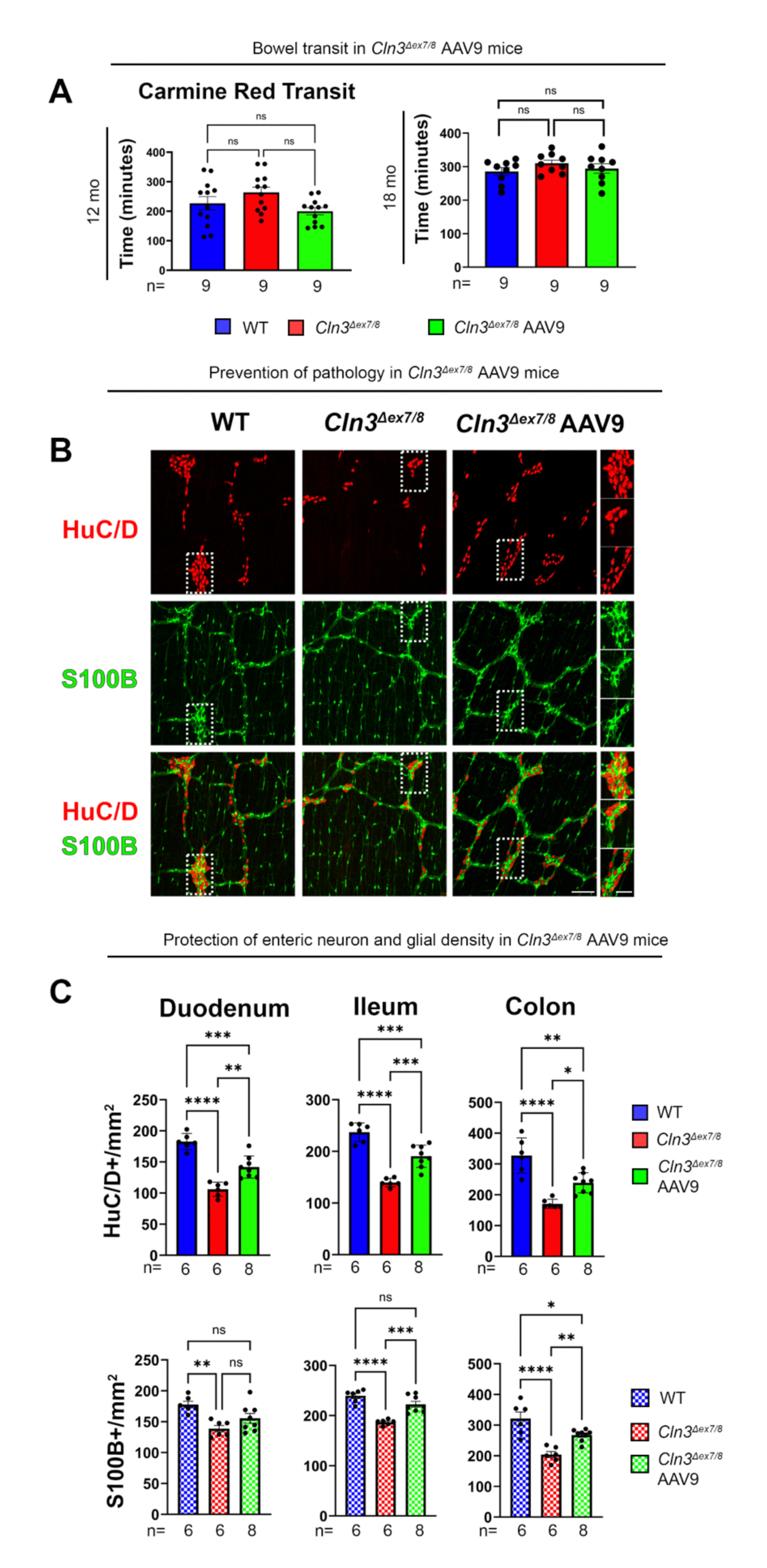
Effects of gene therapy upon the bowel of *Cln3^Δex7/8^* mice. **(A)** Total bowel transit was compared at 12 and 18 months (mo) in untreated *Cln3^Δex7/8^* mice, *Cln3^Δex7/8^* mice treated with AAV9-hCLN3 as neonates (P1) and age-matched wildtype (WT) controls. This was done by measuring the time for gavaged Carmine red to appear in stool. **(B)** Representative photomicrographs of whole-mount bowel preparations of the ileum immunostained for the pan- neuronal marker HuC/D (red) and S100B (green) reveal the protective effects of neonatal intravenous AAV9-hCLN3 treatment upon myenteric plexus neurons and glia in the bowel of disease endstage *Cln3^Δex7/8^* mice treated at P1 vs. WT controls at 18 months. Scale bars 200µm, 50µm higher magnification inserts. (**C**) Measurements of the density of HuC/D positive neurons and glia reveal more neurons present in all bowel regions of 18-month-old *Cln3^Δex7/8^* mice treated neonatally with AAV9-hCLN3 than in untreated *Cln3^Δex7/8^* mice. One-way ANOVA with a post-hoc Bonferroni correction **(A, C)**, * p≤0.05, ** p≤0.01, *** p≤0.001, **** p≤0.0001. Data ± SEM.

Specifically, in 18-month-old AAV9.hCLN3 treated *Cln3^Δex7/8^* mice, the degree of HuC/D+ neuron loss was approximately half of than that seen in untreated *Cln3^Δex7/8^* mice of the same age, with this effect evident along the entire length of the intestine and colon **(Figure 3C**, **Table 2)**. For example, in the duodenum HuC/D+ neuron loss was to 42% of WT values in untreated *Cln3^Δex7/8^* mice vs. 22% of WT values in AAV9.hCLN3 treated *Cln3^Δex7/8^* mice; in the ileum HuC/D+ neuron loss was to 41% of WT values in untreated *Cln3^Δex7/8^* mice vs. 20% of WT values in AAV9.hCLN3 treated *Cln3^Δex7/8^* mice; in the colon HuC/D+ neuron loss was to 48% of WT values in untreated *Cln3^Δex7/8^* mice vs. 27% of WT values in AAV9.hCLN3 treated *Cln3^Δex7/8^* mice. There was also a protective effect of AAV9.hCLN3 treatment upon the density of S100B+ enteric glia, which was significant in the distal part of the small intestine and in the colon at 18 months.

**Table 2.** *Cln3^Δex7/8^* AAV9 Treated Histology Data at 12 months

| HuC/D | *Cln3*<sup>Δex7/8</sup> AAV9<br>neurons/mm<sup>2</sup> | Percent<br>(%) Loss<br>*Cln3*<sup>Δex7/8</sup><br>AAV9 vs.<br>WT | p-value<br>(AAV9-treated<br>vs. WT) | Percent (%)<br>Loss<br>Untreated<br>*Cln3*<sup>Δex7/8</sup><br>vs. WT | % Change<br>Treated<br>vs.<br>Untreated<br>*Cln3*<sup>Δex7/8</sup> | p-value<br>Treated<br>vs.<br>Untreated<br>*Cln3*<sup>Δex7/8</sup> |
| --- | --- | --- | --- | --- | --- | --- |
| Duodenum | 125.60±3.30 | 19.70% | 0.0071 | 20.91% | 101.53% | >0.9999 |
| Ileum | 176.65±9.36 | 15.73% | 0.1014 | 21.66% | 107.57% | >0.9999 |
| Colon | 359.92±34.64 | 9.28% | >0.9999 | 34.43% | 138.37% | 0.0643 |

| S100B | *Cln3*<sup>Δex7/8</sup> AAV9<br>Ganglionic glia/<br>mm<sup>2</sup> | Percent<br>(%) Loss<br>*Cln3*<sup>Δex7/8</sup><br>AAV9 vs.<br>WT | p-value<br>(AAV9-treated<br>vs. WT) | Percent (%)<br>Loss<br>Untreated<br>*Cln3*<sup>Δex7/8</sup><br>vs. WT | % Change<br>Treated<br>vs.<br>Untreated<br>*Cln3*<sup>Δex7/8</sup> | p-value<br>Treated<br>vs.<br>Untreated<br>*Cln3*<sup>Δex7/8</sup> |
| --- | --- | --- | --- | --- | --- | --- |
| Duodenum | 152.19±2.27 | 9.27% | 0.2954 | 8.14% | 98.77% | >0.9999 |
| Ileum | 213.65±8.34 | -0.76% | >0.9999 | 13.38% | 116.32% | 0.1111 |
| Colon | 397.00±14.38 | -1.53% | >0.9999 | 9.00% | 111.57% | 0.6174 |

***Cln3*<sup>Δex7/8</sup> AAV9 Treated Histology Data at 18 months**
| HuC/D | *Cln3*<sup>Δex7/8</sup> AAV9<br>neurons/mm<sup>2</sup> | Percent<br>(%) Loss<br>*Cln3*<sup>Δex7/8</sup><br>AAV9<br>vs. WT | p-value<br>(AAV9-treated<br>vs. WT) | Percent (%)<br>Loss<br>Untreated<br>*Cln3*<sup>Δex7/8</sup><br>vs. WT | % Change<br>Treated<br>vs.<br>Untreated<br>*Cln3*<sup>Δex7/8</sup> | p-value<br>Treated<br>vs.<br>Untreated<br>*Cln3*<sup>Δex7/8</sup> |
| --- | --- | --- | --- | --- | --- | --- |
| Duodenum | 141.86±6.31 | 22.31% | 0.0003 | 41.87% | 133.64% | 0.001 |
| Ileum | 190.53±7.64 | 19.62% | 0.0004 | 41.07% | 136.40% | 0.0002 |
| Colon | 238.85±11.57 | 27.06% | 0.0015 | 47.84% | 139.83% | 0.013 |

| S100B | *Cln3*<sup>Δex7/8</sup> AAV9<br>Ganglionic glia/<br>mm<sup>2</sup> | Percent<br>(%) Loss<br>*Cln3*<sup>Δex7/8</sup><br>AAV9 vs.<br>WT | p-value<br>(AAV9-treated<br>vs. WT) | Percent (%)<br>Loss<br>Untreated<br>*Cln3*<sup>Δex7/8</sup><br>vs. WT | % Change<br>Treated<br>vs.<br>Untreated<br>*Cln3*<sup>Δex7/8</sup> | p-value<br>Treated<br>vs.<br>Untreated<br>*Cln3*<sup>Δex7/8</sup> |
| --- | --- | --- | --- | --- | --- | --- |
| Duodenum | 155.56±7.86 | 17.44% | 0.0906 | 21.99% | 105.83% | 0.2676 |
| Ileum | 222.49±5.88 | 12.45% | 0.0894 | 21.96% | 112.19% | 0.0003 |
| Colon | 267.51±6.87 | 18.97% | 0.0273 | 36.53% | 127.67% | 0.0084 |

## DISCUSSION

The debilitating GI symptoms that occur in many pediatric neurodegenerative disorders are attributed to a variety of causes including CNS degeneration, loss of extrinsic bowel innervation. lack of ambulation, poor hydration, loss of voluntary muscle coordination needed to efficiently pass stool, and effects of medications. Our data from both human and murine CLN3 disease provide new evidence for direct effects of CLN3 disease upon the bowel. This includes degeneration of enteric neurons and glia (this study), and a pathological impact upon bowel smooth muscle^62^. These novel pathologies may underlie or contribute to the debilitating gastrointestinal problems of children with CLN3 disease, revealing new therapeutic targets. Our findings highlight the need to treat the enteric nervous system of the bowel in neurodegenerative disorders, since ENS disease is unlikely to be effectively treated by CNS-directed therapies and ENS disease can be treated by bowel-directed gene therapy, at least in mice, as our data show.

CLN3 disease exerts devastating neurological and neurodegenerative effects upon the CNS resulting in visual impairment, seizures, and cognitive deficits, ending in a premature death in the second or third decade of life^16–18,31^. For this reason, preclinical studies and experimental therapies have understandably focused on treating the CNS via gene therapy or small molecule drugs that target the downstream effects of CLN3 deficiency^16–21,38–40,55–61,64,65^. Although promising effects of CNS-directed gene therapy have been reported in mice^38–40^, and a clinical trial of this strategy was initiated (https://clinicaltrials.gov/study/NCT03770572), none these efforts have proved effective. This may in part be due to incompletely treating the CNS via gene therapy, but also hints at untreated pathology in the rest of the body, a topic that is an active area of investigation.

Such CNS-directed studies are based on the clinical presentation of human CLN3 disease, which includes a broad range of symptoms that may be secondary to CNS dysfunction. GI symptoms include constipation, bowel distension and abdominal pain and discomfort, which are anecdotally recognized although, as yet, very little published information exists about these symptoms. In addition to GI symptoms, CLN3 disease manifestations include cardiac arrhythmia, autonomic dysfunction and peripheral sensory abnormalities, clumsiness, slow or dysregulated movements, and subsequent loss of ambulation and muscular atrophy^16–18,31,66–68^. We hypothesized that these symptoms may be due at least in part to disease outside the CNS. Consistent with this hypothesis, we recently described progressive neuromuscular pathology and myopathy in CLN3 disease^62^, with denervation of the neuromuscular junction (NMJ), loss of NMJ terminal Schwann cells, and marked atrophy of skeletal myofibers in *Cln3^Δex7/8^* mice^62^. Similar pathology was apparent in an early-stage human CLN3 skeletal muscle biopsy^62^.

Now we provide robust evidence that the ENS is affected by CLN3 deficiency. Our data reveal profound pathological effects upon enteric neurons and glia in human CLN3 small bowel and colon at autopsy, with similar progressive enteric pathology in *Cln3^Δex7/8^* mice after a period of apparently normal development. This ENS degeneration may explain the striking dilation in all bowel regions of *Cln3^Δex7/8^* mice and atrophy of bowel smooth muscle layers we reported^62^, since bowel distension is a common feature or both neuropathic and myopathic bowel dysmotility.

Interestingly, when all enteric neurons are missing, the bowel tonically contracts, as demonstrated by human Hirschsprung disease^69^, suggesting the *Cln3^Δex7/8^* mouse bowel phenotype results from an imbalance in neuron subtypes, dysfunction of neurons present, or inability to properly coordinate propagating bowel motility due to patchy degeneration of the ENS. These questions are topics for future study.

These new results are consistent with archival case reports briefly mentioning storage material in myenteric plexus neurons in multiple forms of NCL, which permitted use of rectal biopsy as an NCL diagnostic test^70–73^. These early case reports did not, however, identify neurodegeneration in the ENS, which is quite striking in the human CLN3 disease bowel imaged here at autopsy. This human CLN3 ENS pathology closely resembles ENS pathology in human CLN1 disease at autopsy that we recently described^41^. In addition, *Cln3^Δex7/8^* mouse ENS pathology appears similar to ENS pathology in CLN1 and CLN2 mice that model earlier onset forms of NCL. Like the *Cln3^Δex7/8^* mice, CLN1 and CLN2 disease mouse models display progressive enteric neuron degeneration after a period of apparently normal ENS development, but CLN1 and CLN2 mice have more pronounced bowel transit defects^41^. One unusual feature of all of these lysosomal storage disease models (CLN1, CLN2, and CLN3 disease) is patchy ENS degeneration.

Specifically, in these models, some ENS ganglia appear to be completely missing or markedly damaged, while other nearby ganglia appear fairly normal^41^. To the best of our knowledge, this type of patchy ENS degeneration has not been reported in other mouse models, and suggests degeneration of one ENS cell accelerates degeneration of nearby ENS cells in these CLN models.

A distinguishing feature of ENS pathology in *Cln3^Δex7/8^* mice is that enteric glia are affected to a greater extent than in either CLN1 or CLN2 mice^41^. This may hint at different cellular mechanisms in these lysosomal neurodegenerative diseases, as is becoming evident in the CNS^31,64,65^. Studies are underway to address the underlying molecular mechanisms and the cellular specificity of these effects upon enteric neurons and glia. Nevertheless, our data collectively show ENS disease and bowel dysmotility or distention is present in multiple forms of NCL, and it will be important to assess ENS pathology and bowel function in other similar disorders.

## Clinical implications

Our data provide novel evidence for a pathological impact upon enteric neurons and glia in human CLN3 disease, which is recapitulated in *Cln3^Δex7/8^* mice that have distended bowel and smooth muscle atrophy^62^. The similarity between human and murine data are important because human pathology for bowel collected at autopsy might reflect both disease-associated tissue injury and post-mortem tissue preservation artifacts. From a treatment perspective, CLN3 is a transmembrane protein^22^, so gene therapy is predicted to treat only cells that are transduced.

This means that these newly identified bowel pathologies are very unlikely to be treated by CNS-directed gene therapy. This failure to treat CLN3 disease outside the CNS may explain the why CNS-directed therapies are incompletely effective in *Cln3^Δex7/8^* mice^38–40^. Indeed, it is likely that both CNS-directed and systemic gene therapy will be required to effectively treat well-established CNS phenotypes^26–31^, along with neuromuscular and peripheral nervous system pathology we recently described^62^ and disease effects upon the bowel reported here. One strategy to prevent degeneration of the CLN3 disease ENS is intravenous AAV-mediated gene therapy, as we show. These data extend our recent observations in CLN1 and CLN2 disease mice, where gene therapy also prevented bowel transit defects and extended lifespan^41^. The promising data in CLN1 and CLN2 disease mice were achieved by transducing only a relatively small (15-20%) proportion of enteric neurons^41^, but in these diseases, AAV-expressed lysosomal enzymes are secreted from transduced cells to be taken up by adjacent cells to ‘cross-correct the enzyme deficiencies. That we achieved similar rescue of enteric neurons in *Cln3^Δex7/8^* mice where the transmembrane CLN3 protein is not secreted, is encouraging. We also described positive treatment effects of intravenous AAV9.hCLN3 upon enteric glia and bowel smooth muscle^62^, neither of which are transduced by this virus^41^. Since the CLN3 protein cannot be secreted to cross-correct adjacent cells, this suggests that glial and smooth muscle defects may occur primarily as a result of CLN3 disease-induced enteric neuronal dysfunction. Nevertheless, there may be scope for further improvement using novel viral capsids that transduce a higher proportion of neurons or cell-type specific promoters to drive hCLN3 expression in other affected cell populations. Such targeted strategies may also be required to minimize the risk of hepatocellular carcinoma, which we observed in 6 out of 10 of the AAV9.hCLN3 treated *Cln3^Δex7/8^* mice. Hepatocellular carcinoma is a well-recognized complication of such relatively high doses intravenous AAV9 gene therapy^74,75^, and strategies to mitigate this risk are an active area of investigation.

## Supporting information

Table of Antibodies

## Acknowledgments

This study was initiated because of discussions with affected Batten disease families about their gastrointestinal symptoms. We thank these families for sharing their experiences and inspiring this work. We would particularly wish to thank the family of the child with CLN3 disease who generously donated human tissue samples for our research, and thank all involved in the collection and preservation of these samples. The ANNA-1 anti-HuC/D antibody was kind gift of Dr. Vanda A Lennon, Mayo Clinic, Rochester, MN. We thank Drs. Tom Gillingwater, Tom Wishart and Alison Barnwell for constructive comments on the manuscript. We also thank Kamil Ziółkowski for invaluable assistance in graphing data.

## Study Approvals

All animal procedures were performed in accordance with National Institutes of Health (NIH) guidelines under protocols 2018-0215 and 21-0292 approved by the Institutional Animal Care and Use Committee (IACUC) at Washington University School of Medicine in St. Louis, MO; and The Children’s Hospital of Philadelphia IACUC protocol IAC 22-001041. The analysis of de-identified human autopsy material was reviewed by the Institutional Review Board of Washington University School of Medicine in St. Louis, MO and was considered not to meet the definition of human research and therefore exempt from IRB approval. Organ donor colon was from the Gift of Life Donor Program (also Institutional Review Board exempt).

## Abbreviations

AAV9: Adeno-associated virus 9
AFSM: Autofluorescent storage material
ANOVA: Analysis of variants
BABB: Benzyl alcohol-benzyl benzoate
CLN1 disease: Neuronal ceroid lipofuscinosis 1
CLN2 disease: Neuronal ceroid lipofuscinosis 2
CLN3 disease: Neuronal ceroid lipofuscinosis 3
CNS: Central nervous system
DAPI: 4′,6-diamidino-2-phenylindole
ENS: Enteric nervous system
FITC: Fluorescein isothiocyanate
GFAP: Glial fibrillary acidic protein
GI: Gastrointestinal
IAUCC: Institutional Animal Care and Use Committee
JNCL: Juvenile neuronal ceroid lipofuscinosis
NCL: Neuronal ceroid lipofuscinoses
SCMAS: Subunit c of mitochondrial ATP synthase
PBS: Phosphate buffered saline
TBS: Tris buffered saline
TBST: Tris buffered saline with Triton X-100

## Conflict of interes

JDC has received research support from BioMarin Pharmaceutical Inc., Abeona Therapeutics Inc., REGENXBIO Inc. and Neurogene, and is a consultant for JCR Pharmaceuticals. ROH was a consultant for BlueRock Therapeutics, served on a Scientific Advisory Board for Takeda, and serves on a Scientific Advisory Board for Neurenati Therapeutics. The remaining authors declare no conflicts of interest.

## Authors’ contributions

JDC, ROH, MSS and EAZ conceived and designed the study; JDC, ROH, and MSS obtained funding; EAZ, MJJ, JDC, and SHW carried out the mouse bowel preparations; EAZ performed bowel quantitative analyses; EAZ prepared all the figures with JDC; LLW performed bowel transit studies; EME and LLW managed the mouse colonies, generated the mice for these studies and performed all genotyping; MS and JPS constructed the AAV9-CLN3 vector used in these studies; ITW made the arrangements for donation and collection of the human CLN3 bowel, liaising with the family and health care facility to arrange informed consent; RPB performed the human bowel wholemount immunostaining and imaging under the supervision of ROH; MSS and LLW performed the neonatal vector administration; JDC, MSS and ROH supervised all studies; JDC, MSS, ROH and EAZ interpreted data; JDC and EAZ wrote the manuscript with input from all the authors. All authors read and approved the final manuscript.

## Data Transparency Statement

All the data from this study will be made available to other researchers or is contained in a supplementary data file.

---

**Figure S1.**
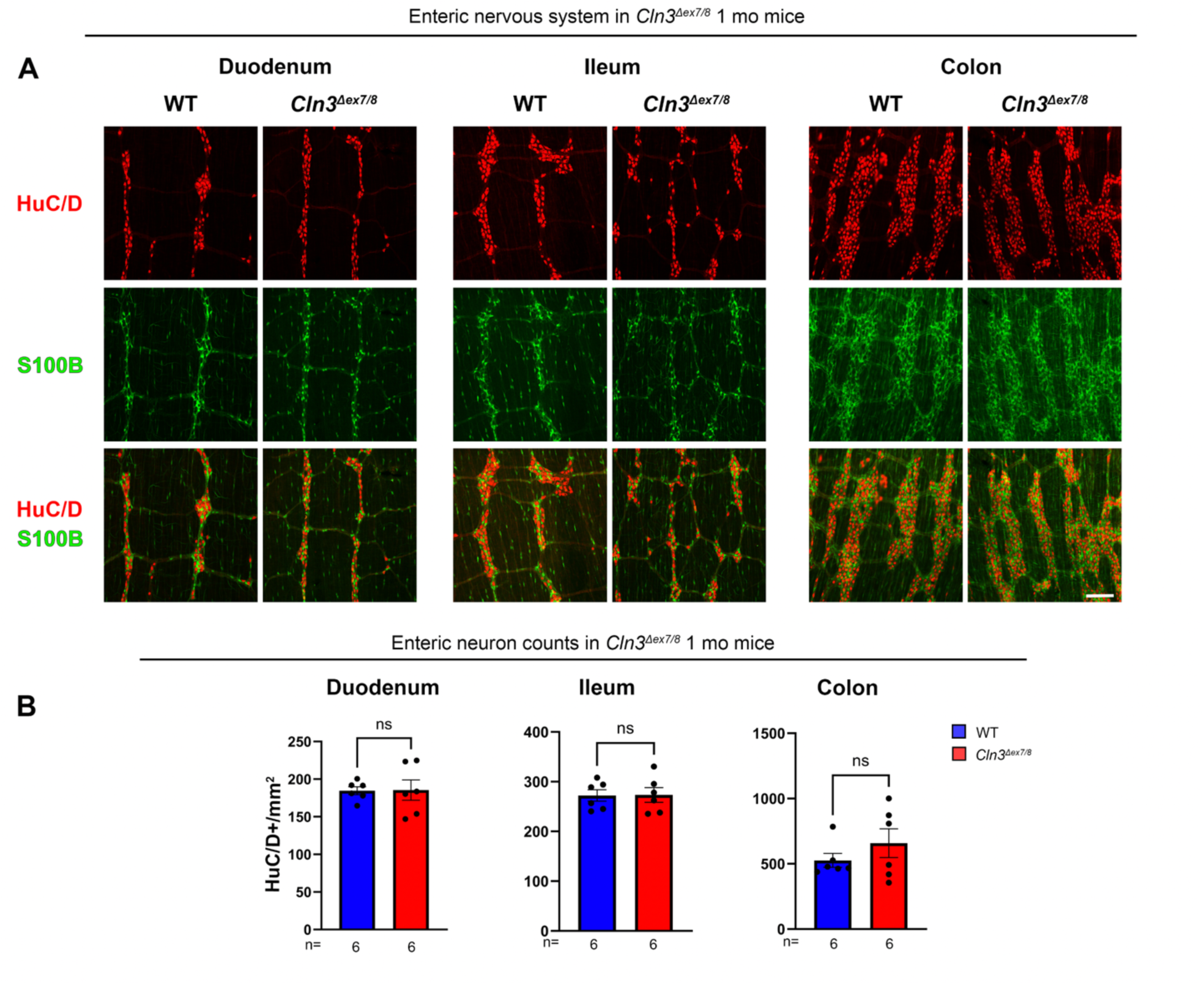
Lack of readily detectable enteric nervous system pathology at 1 month of age. (A) Immunostaining for HuC/D (neurons, red) and the glial marker S100B (green) reveal the myenteric plexus of 1 month old *Cln3^Δex7/8^* mice appears similar to age matched WT mice in duodenum, ileum and colon. Scale bar 200µm. **(B)** Counts of the density of HuC/D+ neurons reveal no significant difference in any bowel regions of 1 month old *Cln3^Δex7/8^* mice compared to age matched WT controls. Unpaired t-tests **(B)**. Data ± SEM.

**Figure S2.**
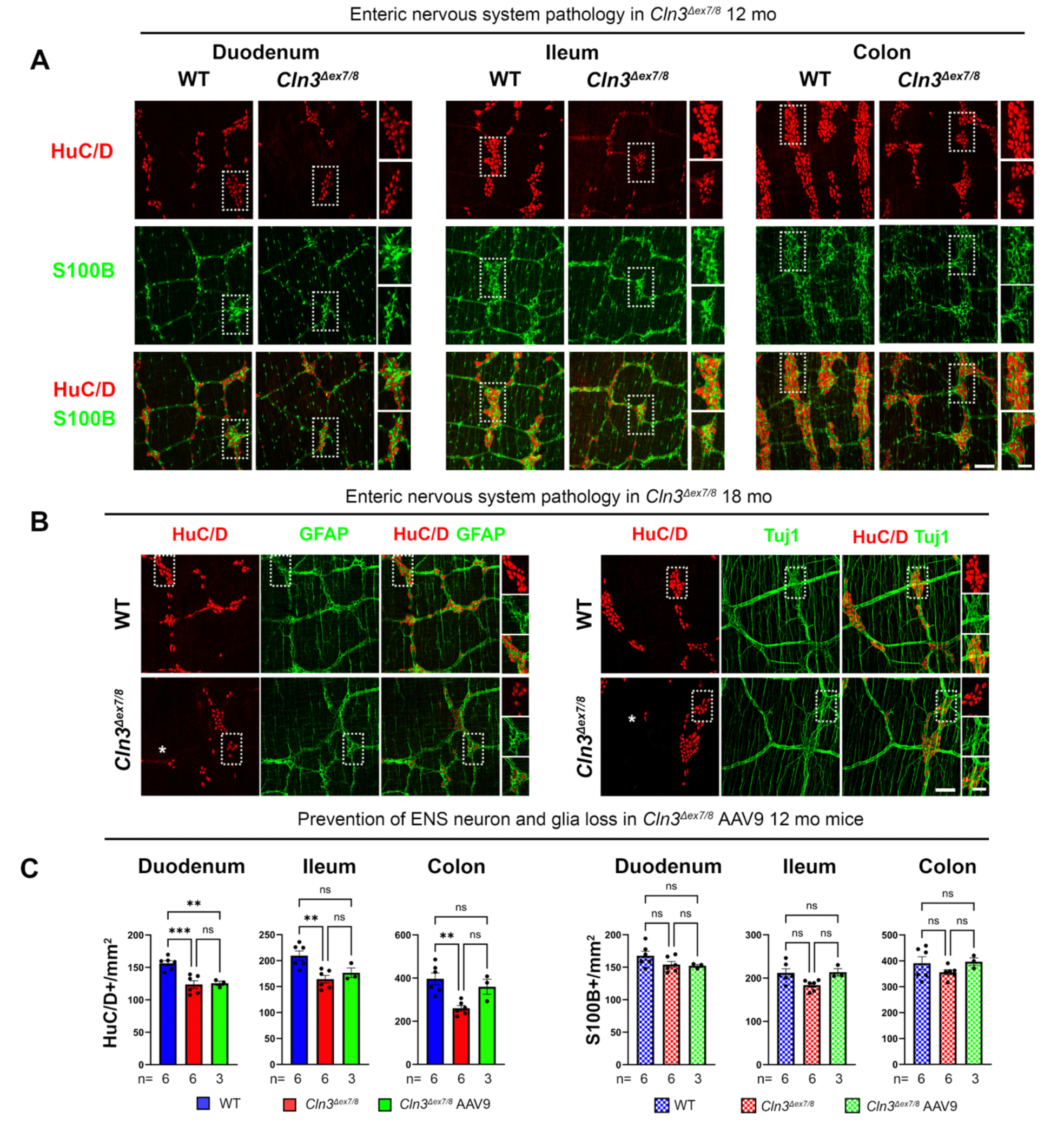
Evidence for enteric nervous system pathology in *Cln3^Δex7/8^* mice at 12 and 18 months. **(A)** Photomicrographs show immunostaining for HuC/D (neurons, red) and the glial marker S100B (green) in the duodenum, ileum and colon *Cln3^Δex7/8^* mice and age-matched wildtype (WT) controls at 12 months of age (mo). Scale bars 200µm, 50µm in higher magnification views. **(B)** Photomicrographs showing immunostaining for HuC/D (neurons, red) and either the glial marker glial fibrillary acidic protein (GFAP, green) or the neuronal microtubule marker Tuj1 (green) in the duodenum *Cln3^Δex7/8^* mice and age-matched WT controls at 18 months of age. Scale bars 200µm, 50µm in higher magnification views. Asterix (*) indicates regions of near complete enteric neuron loss. **(C)** Measurements of the density of HuC/D+ myenteric neurons and S100B+ myenteric glia reveal no significant differences (ns) in all bowel regions of 12-month-old *Cln3^Δex7/8^* mice treated with AAV9-hCLN3 vs. untreated *Cln3^Δex7/8^* mice. Data ± SEM. One-way ANOVA with a post-hoc Bonferroni correction, ** p≤0.01, *** p≤0.001.

