## Supplementary material for "Enteric nervous system degeneration in human and murine CLN3 disease, is ameliorated by gene therapy in mice": Table of Antibodies

| **Antibody** | **Concentration** | **Catalog number** | **Source** |
| --- | --- | --- | --- |
| ANNA-1 (HuC/D) | 1:10000 (mouse bowel) | N/A | Kind gift from Dr. Vanda Lenon, Mayo Clinic; RRID: AB_2314657 |
| Mouse anti HuC/D | 1:200 (human bowel) | A21271 | Invitrogen  RRID: AB_221448 |
| Rabbit anti-SOX10 | 1:100 (human bowel) | 383R-15 | Cell Marque  RRID: AB2941085 |
| Rabbit anti-GFAP | 1:10000 (mouse bowel) | Z0334 | Agilent  RRID: AB_10013382 |
| Rabbit anti-beta III tubulin (Tuj1) | 1:5000 (mouse bowel) | ab18207 | Abcam  RRID: AB_444319 |
| Chicken anti-beta III tubulin (Tuj1) | 1:500 (human bowel) | GTX85469 | GeneTex  RRID: AB_10629222 |
| Rabbit anti-S100 beta | 1:300 (mouse bowel) | ab52642 | Abcam  RRID: AB_882426 |
| AlexaFluor goat anti-human 546 | 1:400 (mouse bowel) | A-21089 | ThermoFisher Scientific (Invitrogen)  RRID: AB_2535745 |
| AlexaFluor goat anti-rabbit 488 | 1:400 (mouse bowel) | A-11008 | ThermoFisher Scientific (Invitrogen)  RRID: AB_143165 |
| AlexaFluor donkey-anti mouse 594 | 1:400 (human bowel) | A-21203 | ThermoFisher Scientific (Invitrogen)  RRID: AB_141633 |
| AlexaFluor goat-anti chicken 594 | 1:400 (human bowel) | A-21449 | ThermoFisher Scientific (Invitrogen)  RRID: AB_2535866 |

**Table S1 Antibodies used**
